# Mice with humanized livers reveal the involvement of hepatocyte circadian clocks in rhythmic behavior and physiology

**DOI:** 10.1101/2022.09.08.506890

**Authors:** Anne-Sophie Delbès, Mar Quiñones, Cédric Gobet, Julien Castel, Raphaël G. P Denis, Jérémy Berthelet, Benjamin D. Weger, Etienne Challet, Aline Charpagne, Sylviane Metairon, Julie Piccand, Marine Kraus, Bettina H. Rohde, John Bial, Elizabeth M. Wilson, Lise-Lotte Vedin, Mirko E. Minniti, Matteo Pedrelli, Paolo Parini, Frédéric Gachon, Serge Luquet

## Abstract

The circadian clock is an evolutionarily acquired gene network that synchronizes physiological processes to adapt homeostasis to the succession of day and night. While most mammalian cells have a circadian clock, their synchronization at the body-level depends on a central pacemaker located in the suprachiasmatic nuclei of the hypothalamus that integrates light signals. However, peripheral organs are also synchronized by feeding cues that can uncoupled them from the central pacemaker. Nevertheless, the potential feedback of peripheral signals on the central clock remains poorly characterized. To discover whether peripheral organ circadian clocks may affect the central pacemaker, we used a chimeric model in which mouse hepatocytes were replaced by human hepatocytes. These human hepatocytes showed a specific rhythmic physiology caused by their blunted response to mouse systemic signals. Strikingly, mouse liver humanization reprogrammed the liver diurnal gene expression and modified the phase of the circadian clock. The phase advance was also reflected in the muscle as well as the entire rhythmic physiology of the animals, indicating an impact on the circadian function of the central clock. Like mice with a deficient central clock, the humanized animals shifted their rhythmic physiology more rapidly to the light phase under day feeding. Our results indicate that peripheral clocks may affect the central pacemaker and offer new perspectives to understand the impact of peripheral clocks on the global circadian physiology.

## INTRODUCTION

To adapt homeostasis to the changing environment present on Earth, organisms from bacteria to mammals have evolved a timing system that anticipates these changes. This endogenous timing system, called the circadian clock, orchestrates most aspects of physiology and behavior. The mammalian circadian system is hierarchically organized. A central clock localized in the suprachiasmatic nucleus (SCN) of the hypothalamus is daily synchronized by light via the retino-hypothalamic tract and coordinates the peripheral clocks localized in peripheral tissues (Hastings et al., 2018). The SCN synchronizes most aspects of circadian physiology and is required to keep phase coherence between the different peripheral organs (Ralph et al., 1990; Sinturel et al., 2021; Yoo et al., 2004). Nonetheless, how the SCN orchestrates this phase coherence is still only partially understood.

However, in the absence of SCN, the individual peripheral organs continue to show robust rhythms, indicating the existence of self-sustained cellular clocks (Sinturel et al., 2021; Yoo et al., 2004). In all mammalian cells, this rhythmicity is generated by a molecular machinery consisting of interconnected transcriptional and translational feedback loops in which the transcription factor Brain and Muscle Aryl hydrocarbon receptor nuclear translocator-Like 1 (BMAL1, encoded by the *Arntl* gene) plays a critical role (Patke et al., 2020). Abolishing BMAL1 activity in mouse hepatocytes demonstrated that around 35% of liver rhythmic genes depend on a functional circadian clock whereas the other rhythmic genes are regulated by systemic and/or feeding cues (Johnson et al., 2014; Kornmann et al., 2007; Weger et al., 2021). Nevertheless, disrupting hepatocyte clock has no impact on the central clock in the SCN (Johnson et al., 2014; Lamia et al., 2008). On the other hand, restoring *Bmal1* expression and a functional circadian clock in a full-body *Bmal1* knockout (KO) animal has no effect on the global rhythmic activity and behavior of these animals (Koronowski et al., 2019). Therefore, while the central clock synchronizes peripheral clocks, there is so far no evidence that peripheral clocks can impact the central clock in the SCN.

Also, recent evidence further suggests that the circadian clocks of different cell types communicate between each other inside the same organ. For example, astrocytes circadian clock can entrain neuronal clock in the SCN (Brancaccio et al., 2019; Patton et al., 2022) and is important to determine the circadian period (Barca-Mayo et al., 2017; Tso et al., 2017). Moreover, the disruption of the circadian clock in hepatocytes modulates the rhythmic gene expression of other liver cell types (Guan et al., 2020) and even the synchronization of the circadian clock in other tissues in response to feeding cues (Manella et al., 2021). These studies highlight the importance of the communication between the clocks in the different cell types inside the same tissue or in other more distant tissues (Koronowski and Sassone-Corsi, 2021).

To decipher further the impact of these cellular communications in the organization of liver circadian physiology, we implanted in mouse liver human hepatocytes that are hypothesized to show different physiological properties and responses to systemic and feeding cues compared to mouse hepatocytes (Azuma et al., 2007; Jiang et al., 2020), as well as different rhythmic functions as shown in non-human primates (Mure et al., 2018). Here we show that implantation of human hepatocytes in mice (Liver Humanized Mice: LHM) result in a global loss of rhythmicity in liver gene expression in comparison to control animals implanted with mouse hepatocytes (Liver “Murinized” Mice: LMM). While most circadian clock genes were still rhythmic in the liver of LHM, they displayed an advanced phase and lower amplitude in both mouse and human hepatocytes, suggesting that human hepatocytes can entrain the remaining mouse hepatocytes in the liver of LHM. Strikingly, we discovered that these human hepatocytes also regulate rhythmic gene in the muscle and impact global rhythmic behavior and metabolism, suggesting potential feedback of the implanted human hepatocytes on the hypothalamus and the central clock in the SCN. Finally, during imposed day feeding, LHM adapted more quickly to the light phase, confirming that the engrafed human hepatocytes impacts the entrainment property of the central circadian pacemaker. Altogether, the chimeric LHM provide a yet uncharacterized mechanism by which perturbation of the liver clock might directly alter global circadian physiology.

## RESULTS

### Human hepatocytes in a rodent environment impose specific rhythmic gene expression to surrounding liver cells

It is now established that most cells in the body contain self-sustained molecular clocks that can be synchronized by both signals from the central clock in the SCN and systemic cues including feeding rhythm (Atger et al., 2017; Koronowski and Sassone-Corsi, 2021). Recently, it has been shown that the disruption of the circadian clock specifically in hepatocyte (Guan et al., 2020; Manella et al., 2021) or the presence of a functional clock only in hepatocyte in a clock-depleted animal (Greco et al., 2021; Koronowski et al., 2019) modulate the rhythmic expression in other cell types inside the same tissue or in other tissues, as well as their response to feeding. However, while transplanting SCN cells with different circadian properties in an intact animal can transfer the donor’s SCN circadian properties and impact animal behavior (Ralph et al., 1990), there is no description of such effects for peripheral tissues.

To answer this question, we used chimeric LHM in which mice livers are repopulated with human hepatocytes that are expected to have their own circadian properties and capacities to respond to systemic signals, as described for human and another diurnal primate (Jiang et al., 2020; Mure et al., 2018). In this study we used the FRG^®^-triple KO mouse model that exhibits a mutated Fumarylacetoacetate Hydrolase (*Fah*) and severe immunodeficiency (*Rag2*^*-/-*^ and *Il2rg*^*-/-*^) (Azuma et al., 2007). After transplantation with primary human hepatocytes, the engrafted cells can repopulate the liver up to 80% due to positive selection over rodent hepatocyte functionally impaired for FAH (Azuma et al., 2007). Further improvement of this model was obtained through backcrossing FRG^®^-KO mice with the non-obese diabetic (NOD) mice, which can more easily accept xenografts (Foquet et al., 2017; Takenaka et al., 2007). The resulting model FRGN mice were engrafted with human hepatocytes from a donor or murine hepatocyte from NOD mice to produce respectively liver-humanized (Hu-FRGN®, LHM) or control “murinized” mice (Mu-FRGN®, LMM) (Minniti et al., 2020) (**Figure 1A**). Circulating levels of human albumin (hAlb) in LHM were used to corroborate the extent of human hepatocyte repopulation and were in average 6414±412 µg/ml in LHM while no hAlb could be detected in LMM (**Figure S1A**). Over the course of the entire study, only mice with circulating levels of human albumin >3500 µg/mL (expecting to correspond to human hepatocyte repopulation >70% (Ellis et al., 2013)) were used. An additional functional readout of successful liver humanization was gathered through the measure of triglycerides (TG) content in low density lipoprotein (LDL). While rodents transport TG mainly in high density lipoproteins (HDL) (Ellis et al., 2013), TG in human serum are typically enriched in liver-born LDL. Unlike LMM, LHM lipoprotein profile displayed enrichment of TG in LDL, confirming the extent of liver humanization and in agreement with previous observations (Minniti et al., 2020) (**Figure S1B**).

**Figure 1.**
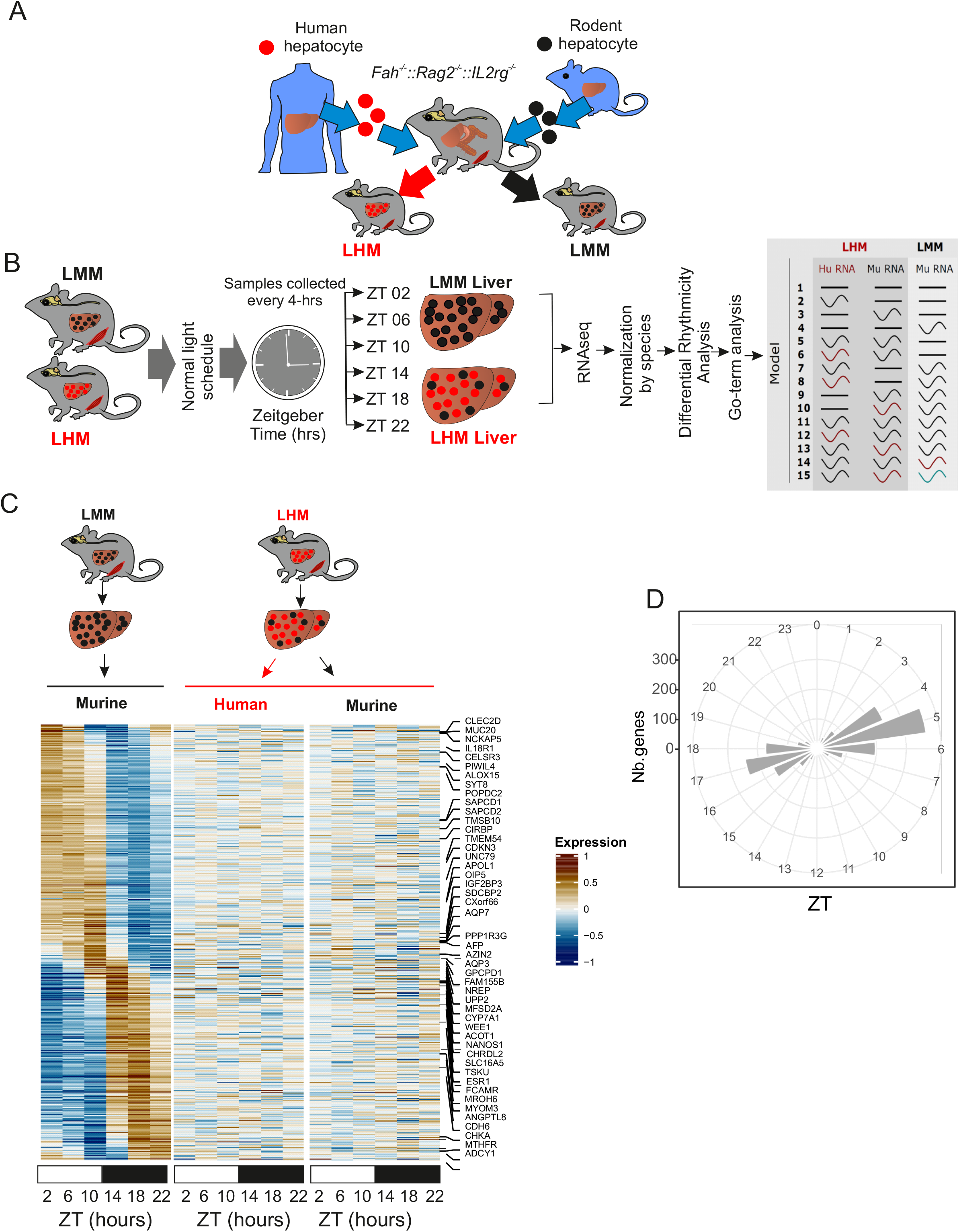
Engraftment of human hepatocyte impacts liver rhythmic gene expression. **A**). Model for humanized *Fah*^-/-^, *Rag2*^-/-^, *Il2rg*^-/-^ (FRG®-KO) that can be repopulated with primary human (red) or murine (black) hepatocytes to produce liver humanized (LHM) or “murinized” (LMM) mice. **B**) Experimental design for liver tissue collection prior to RNA extraction, sequencing and analyzing according to gene expression rhythmic properties. Alteration of rhythmic gene expression of murine transcript in the liver of LMM (black, Mu RNA) and both murine (red, Mu-RNA) and human transcript (red Hu-RNA) in the liver of LHM mice is assessed by model selection (model 1–15): black line, stable transcription; black wave, rhythmic transcription; red or green waves, rhythmic profiles with different rhythmic parameters (i.e., phase and/or amplitude). **C**) Heat maps of normalized rhythmic mRNA levels (BICW > 0.3, log2 amplitude > 0.5) in the liver of LMM (black) and LHM (red) murine and human transcript (red) in the liver of LMM and LHM. RNA presented here belonged to model 4 where genes were rhythmic only in LMM **D**) Radial plot of the distribution peak phase of expression for genes rhythmic only in the liver of LMM (model 4).

To decipher the impact of the transplantation of human hepatocytes on the rhythmic physiology of the mouse liver, we analyzed liver rhythmic gene expression in LMM and LHM. Both LMM and LHM mice, engrafted on the same week and exposed to the same nutritional and pharmacologic treatment (see method) and housing conditions, were sacrificed every 4-hours and liver tissue collected prior to RNA extraction and sequencing. Due to the high depth RNA sequencing, we can discriminate between human and non-human RNA orthologues within LHM liver samples, allowing independent normalization for the two species before the analysis of differential rhythmicity of gene expression using our recently developed *dryR* method (Weger et al., 2021) (**Figure 1B and S1C**). *dryR* assigns groups of genes to various models of rhythmic behavior and species. For instance, genes assigned to model 4 are rhythmic only in LMM while those in models 2, 3, 5, and 6 are arrhythmic in LMM but displayed rhythm in LHM with similar or different species-specific rhythmic parameters (**Figure 1B**).

The analysis reveals that the largest group (model 4) is composed of genes that lose rhythmicity specifically in LHM animals, for both human and mouse transcripts (**Figure 1C, 1D, S1D, Table S1**). These genes show the classical biphasic pattern observed for rhythmic genes regulated by systemic or feeding cues (**Figure 1D**) (Weger et al., 2021). These rhythmic genes are mainly involved in protein synthesis and ribosome biogenesis, suggesting an attenuated activation of the mechanistic target of rapamycin (mTOR) pathway (**Table S2**). A comparison with the genes differentially expressed in the liver of *Raptor* or *Tsc1* KO mice, the main positive and negative regulators of the TORC1 complex, respectively (Bai et al., 2021), confirms that the genes differentially expressed in LHM are indeed enriched in genes transcriptionally regulated by the mTOR pathway (**Figure S1E**). Additional analysis of the phosphorylation of ribosomal protein S6 (RPS6, a bona fide TORC1 target) confirms the absence of the expected rhythmic activation of this pathway in the liver of LHM (Jouffe et al., 2013) (**Figure S1F**). The mTOR pathway being recognized as a key sensor of systemic and nutritional signals (Saxton and Sabatini, 2017), this suggests that the human hepatocytes are unable to respond to such signals when placed in a mouse environment. Alternatively, the human hepatocytes can also respond to these signals in a different manner that does not result in the rhythmic activation of the mTOR pathway. The inappropriate response of human hepatocytes to mouse signals is also illustrated by the blunted response to Growth Hormone (GH), an important regulator of the sex biased liver function and metabolism (Al-Massadi et al., 2022; Lichanska and Waters, 2008). Indeed, endogenous mouse GH is high while IGF-1 is low in LHM (**Figure S1G**). This shows that human hepatocytes cannot secrete IGF-1 in response to GH and, in turn, the low level of IGF-1 is not sufficient to exert its inhibitory action on GH secretion, resulting in constantly high GH levels. As an additional evidence of the disruption of GH signaling, the sex biased gene expression as well as the activation of the sex biased activation of the STAT5 pathway, two recognized readouts of GH signaling (Weger et al., 2019; Zhang et al., 2012), are also perturbed, with an increase of female biased signaling and a decrease of the male biased signaling, likely as a result of the constantly high GH level (**Figure S1H**). The IGF-1 / mTOR pathway being important for the synchronization of the circadian clock by systemic signals (Crosby et al., 2019), this blunted response could contribute to the lack of entrainment of human hepatocytes.

On the other hand, hundreds of genes mainly involved in angiogenesis and telomere maintenance are only rhythmic in human hepatocytes (**model 2, Figure S1D**), suggesting that these cells can express their own specific rhythmic physiological program different from the one of mouse hepatocytes (**Figure S2A, S2B, Table S1, S2**). Of particular interest is model 14 in which 110 genes are found to share similar phase advance in both human and rodent cells in LHM compared to the liver of LMM (**Figure 2A, 2B, Table S1**). Surprisingly, this group shows a clear enrichment to circadian clock related pathways (**Table S2**) and includes most circadian clock and clock target genes (**Figure 2C**). This phase advance of clock genes expression associated with a lower amplitude mirrors the presence of a functional clock only in hepatocytes in a clock depleted animal (Koronowski et al., 2019), suggesting that the human hepatocytes clock in a mouse liver operates like independent oscillators not responding to synchronizing rhythmic systemic signals. To support this hypothesis, we compared our results with the rhythmic genes present in the liver of wild-type, *Bmal1* KO, and *Bmal1* KO with a specific rescue of *Bmal1* only in the liver (liver-RE), an example of independent hepatocyte oscillator in a circadian clock depleted animal (Koronowski et al., 2019). Interestingly, the genes that lost rhythmicity in the liver of LHM (model 4) are specifically enriched in genes rhythmic only in WT animals (**Figure S2C**), supporting the idea that these genes are indeed mainly regulated by systemic signals originating from other organs in a circadian clock-dependent fashion, but independent of the hepatocyte circadian clock. This confirms that the human hepatocytes cannot respond to mouse synchronizing rhythmic systemic signals and operate largely independently from the rest of the organism. The fact that the mouse hepatocytes in LHM mice show the same circadian behavior suggests that the human hepatocytes can entrain the remaining mouse cells in the liver of LHM.

**Figure 2.**
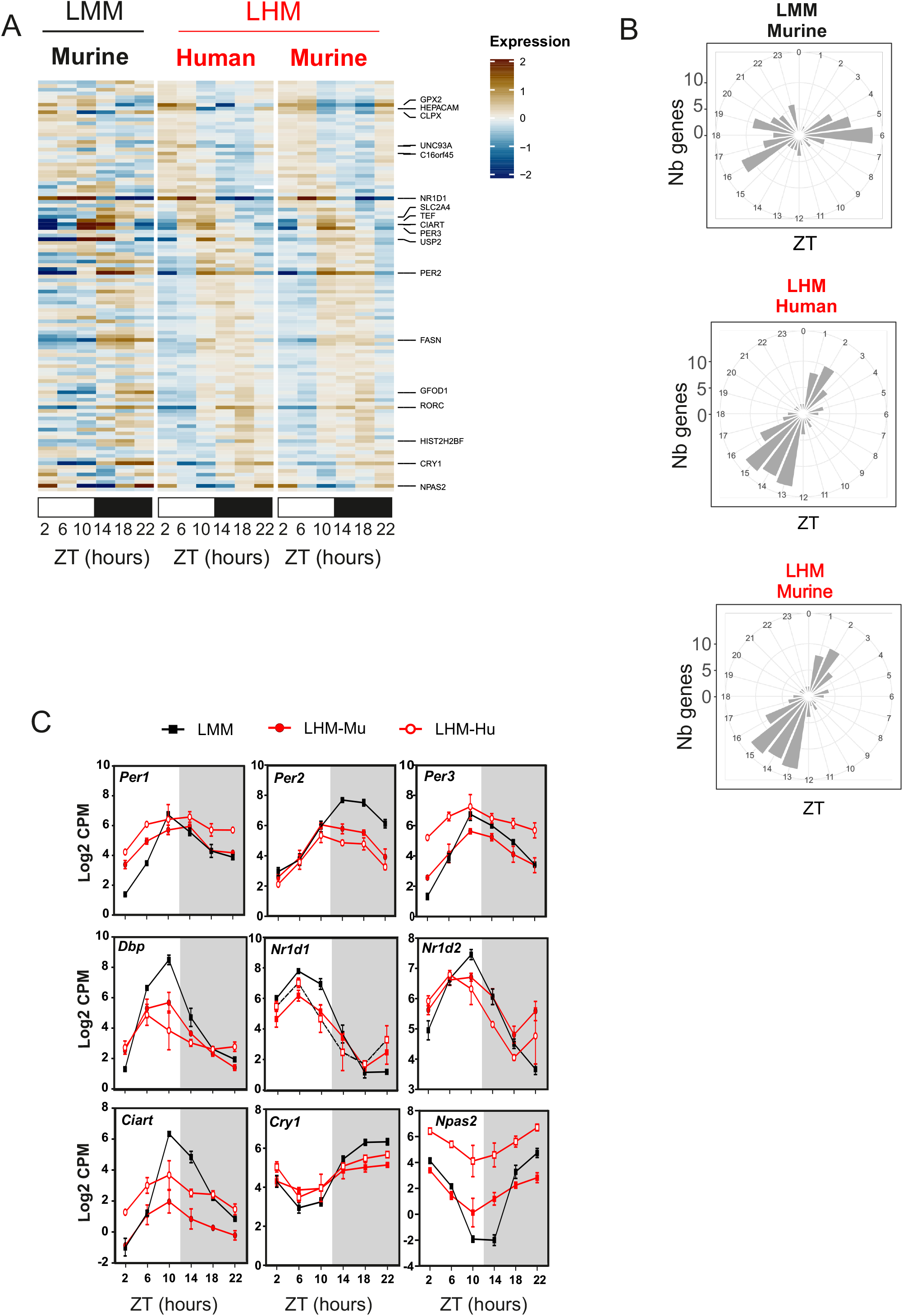
Engraftment of human hepatocyte impacts the phase of the liver circadian clock. **A**) Heat maps of normalized rhythmic mRNA levels (BICW > 0.3, log2 amplitude > 0.5) in the liver of LMM (black) and LHM (red) murine and human transcript (red) in the liver of LMM and LHM. Genes presented here belonged to model 14 where both LHM-Mu & LHM-Hu orthologue transcripts share the same phase which is different from LMM. **B**) Radial plot distribution of the peak phase of expression for rhythmic genes in the liver of LMM and murine and human orthologue in the liver of LHM from model 14. **C**) Rhythmic expression of circadian clock genes for murine (black, LMM) and murine (LHM-Mu) and human (LHM-Hu) RNA orthologue in the humanized liver. Data are expressed as mean +/- SEM (n=12 mice per groups, 2 points per replicates). For statistical details, see Table S1.

### The muscle circadian clock of liver humanized mice is phase advanced

To determine if this entrainment by human hepatocytes is specific to the mouse cells in the liver, we performed a similar RNA-Seq analysis on the skeletal muscle (quadriceps) of LMM and LHM (**Figure 3A**). While the biggest group of genes (model 4: 32%) shows similar rhythms in LHM and LMM, a similar and important proportion of genes (model 2: 26% and model 3: 28%, respectively) are only rhythmic in the muscle of LHM and LMM mice, respectively, showing that human hepatocytes in LHM have a strong influence on the rhythmicity of other tissues (**Figure S3A and Table S1**). The genes that lose rhythmicity in LHM (model 3) are mainly involved in protein lipid and cholesterol metabolisms, showing the impact of the human hepatocytes on the global physiology and metabolism of the animals (**Figure S3B, S3C and Table S3**). Strikingly, the group of genes displaying different circadian parameters (model 5: 13%) includes almost all the circadian clock genes and bona fide clock target genes (**Figure 3B, 3C, S3A, Table S1 and S3**). While we did not find clear effect on the amplitude, all these clock genes show an advanced phase of expression more important than the one we found in the liver (**Figure 3D**). This result demonstrates that the engraftment of human hepatocytes in mouse liver can surprisingly phase advance the circadian clock of other peripheral tissues.

**Figure 3.**
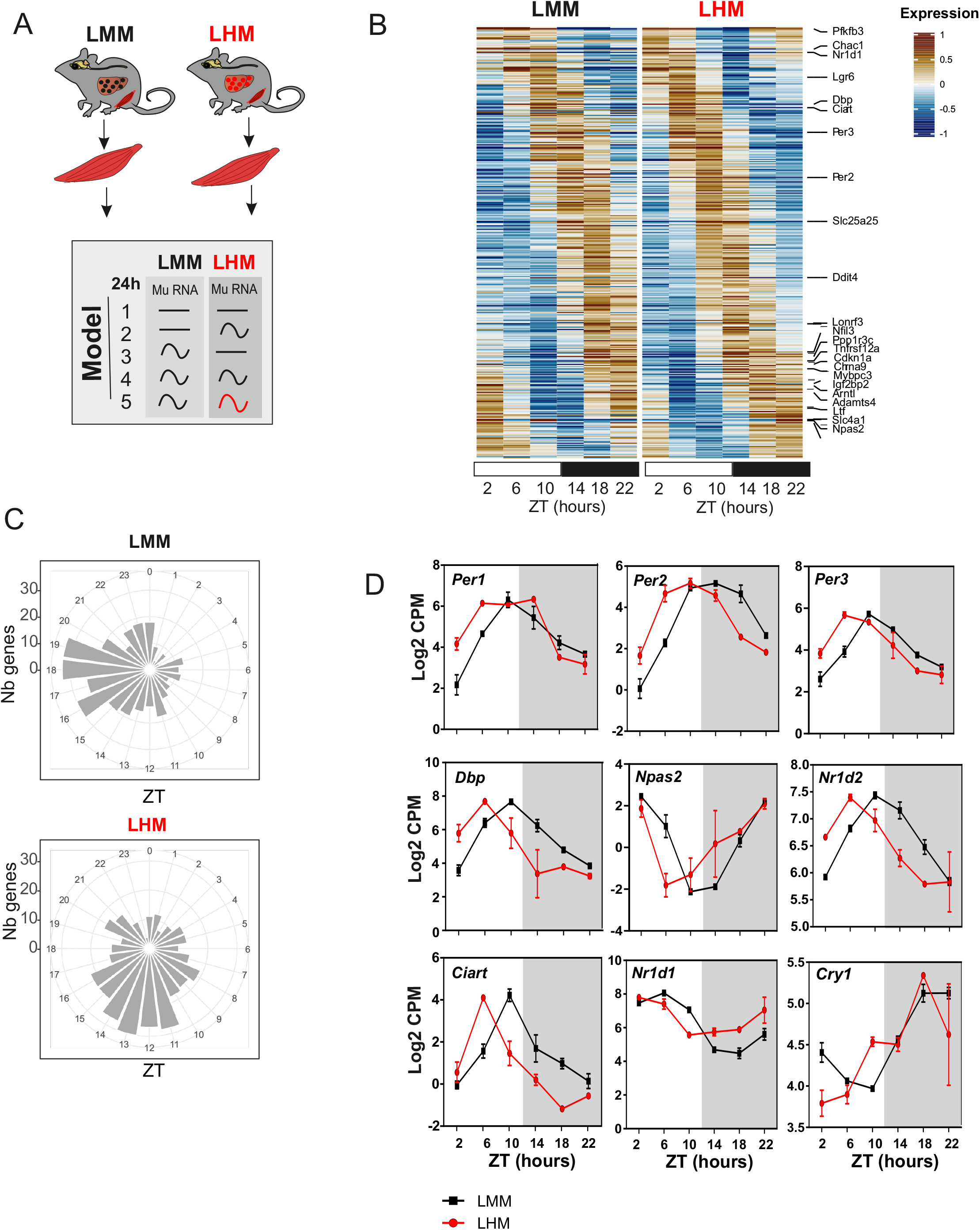
The muscle circadian clock of liver humanized mice is phase advanced. **A**) Experimental design for muscle tissue collection prior to RNA extraction, sequencing and analyzing according to rhythmic properties, alteration of rhythmic gene expression of murine transcript in the muscle of LMM (black) and LHM (red) assessed by model selection (model 1–5): black line, stable transcription; black wave, rhythmic transcription; red wave, rhythmic profiles with different rhythmic parameters (i.e., phase and/or amplitude). **B**) Heat maps of normalized rhythmic muscle mRNA levels (BICW > 0.5, log2 amplitude > 0.5) in LMM (black) and LHM (red) from model 5 where LMM and LHM muscle transcript are rhythmic but with different phase. **C**) Radial plot distribution of the peak phase of expression for rhythmic genes in the liver of LMM and murine and human orthologue in the liver of LHM from model 5. **D**) Rhythmic expression of circadian clock genes in the muscle of LMM and LHM. Data are expressed as mean +/- SEM (n=11-12 animals per groups). For statistical details, see table S1.

### Human hepatocytes advance the phase of circadian metabolism and behavior of liver humanized mice

Because feeding cues act as the main driver of the phase of peripheral clocks, disconnecting them form the entrainment by the SCN (Damiola et al., 2000; Stokkan et al., 2001), we hypothesized that the advanced phase of the muscle circadian clock could be a consequence of a change of rhythmic metabolic and behavioral output driven by the engrafted human hepatocytes. Therefore, we performed a comprehensive characterization of the rhythmic physiology of LMM and LHM. Under a normal light/dark cycle, LHM strikingly display an approximately 2hrs phase advance of locomotor activity, feeding behavior, respiratory exchange ratio, and fat oxidation compared to LMM (**Figure 4A-H**). The amplitude of the rhythmic locomotor activity and food intake is also decreased (**Figure S4A-D**). Of note, LMM and LHM did not significantly differ in average cumulative food intake (**Figure S4B**) and show only a slightly reduced body weight and body fat (**Figure S4E**), suggesting that the change in entrainment properties did not result from metabolic impairment. Altogether, this confirms that the phase advance observed in the muscle of LHM is likely a consequence of the advanced feeding activity.

**Figure 4.**
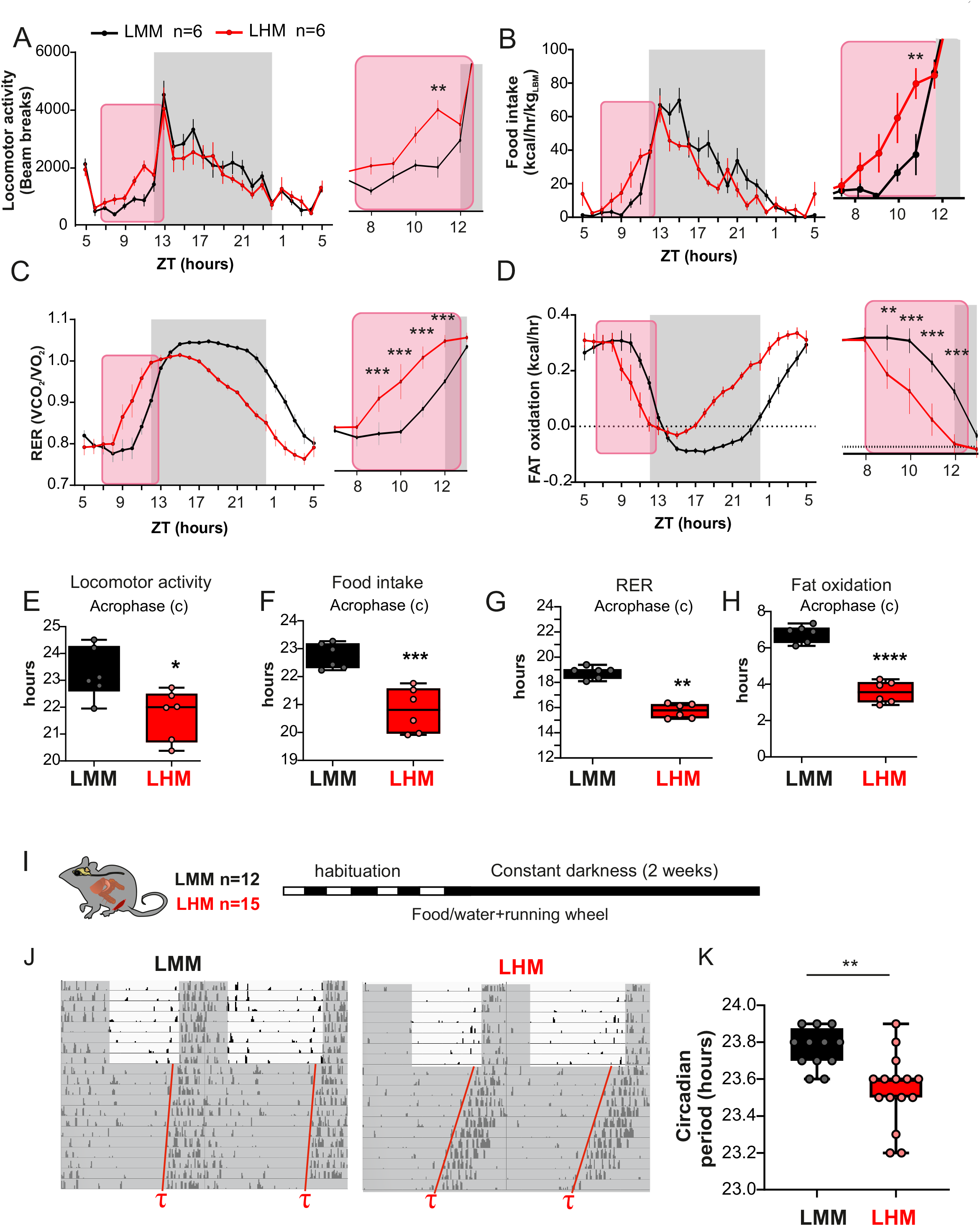
Human hepatocytes advance the phase of circadian metabolism and behavior of liver humanized mice. **A**-**D**) Metabolic evaluation of LMM and LHM represented as 3-days average spontaneous locomotor activity (**A**), food intake (**B**, kcal/hr/kg of lean body weight), respiratory exchange ratio (RER, **C**), and fat oxidation (**D**). Insert are showing magnification of the ZT8-12 time window. **E**-**H**) Cosinor analysis of rhythmic parameters acrophase (c) for locomotor activity (**E**), food intake (**F**), RER (**G**), and fat oxidation (**H**). **I**) Experimental design for the determination the circadian period of LMM and LHM via the recording of the circadian running wheel activity in constant darkness. **J**) Representative actogram and circadian period in LMM and LHM. **K**) Circadian period of locomotor activity in LHM and LMM (n=12 and 15 for LMM and LHM, respectively). Codes for statistical values: * P<0.05, ** P<0.01, *** P<0.0001.

As described in the case of the familial advanced sleep-phase syndrome (FASP), this phase advance observed in light/dark condition could be a consequence of a short circadian period (Hirano et al., 2016; Jones et al., 1999; Xu et al., 2005; Xu et al., 2007). Accordingly, humans with shorter circadian periods show an early chronotype (Brown et al., 2008; Duffy et al., 2001). Thus, we determined the circadian period of the animals via the recording of their circadian wheel-running activity in constant darkness (**Figure I**). Strikingly, LHM show a significantly shorter period (**Figure 4J, 4K**), demonstrating feedback signals from human hepatocytes to the central clock in the SCN. This suggests an unexpected and never described impact of hepatocytes on the circadian function of the SCN.

### Engrafted human hepatocytes impact rhythmic gene expression in the hypothalamus

To decipher how the engrafted human hepatocytes can extend their influence beyond the liver and muscle to the brain centers involved in the control of metabolism and circadian rhythms, we performed a time-resolved RNA-Seq experiment on the hypothalamic brain punches of the Arcuate nucleus (ARC) and the SCN (**Figure 5A**). Relative enrichment of the SCN-enriched gene *Aralkylamine N-Acetyltransferase* (*Aanat*) or ARC-enriched *Agouti related peptide* (*Agrp*) confirms dissection accuracy (**Figure S5A**). The majority (58% and 53%, model 4) of the rhythmic genes keep their rhythmicity in the SCN and the ARC, respectively, including the majority of circadian clock genes (**Figure S5B, S5C and Tables S1**). However, the second largest group of genes representing 30% and 26% of the rhythmic genes (model 3) loses rhythmicity in the SCN and the ARC of LHM, respectively, showing that engraftment of human hepatocytes can unexpectedly impact the rhythmic gene expression in the hypothalamus (**Figures 5B-E, S5B, S5C, and Tables S1**). These genes that lose rhythmicity in the SCN and ARC of LHM are enriched for genes encoding proteins involved in ions transport, critical for the function of neurons (Tables S4 and S5). In addition, a small subset of 2% and 10% of rhythmic genes in the SCN and the ARC, respectively, exhibits different circadian parameters between LMM and LHM (model 5, **Figure 5F-I**). In contrast to liver and muscle, only a few circadian clock genes, including *Nr1d1* and *Dbp*, are included in these groups, both showing a phase advance in LHM animals (**Figure S5D, S5E**). Nevertheless, additional analysis shows that the genes in models 4 and 5 exhibit both an enrichment for genes controlled by the circadian regulator CLOCK (Alhopuro et al., 2010). This suggests that only a subset of circadian clock-controlled genes are phase advanced in the hypothalamus, potentially because of conflicting light signals synchronizing the SCN clock (Shigeyoshi et al., 1997; Xu et al., 2021) (**Figure S5F**). Altogether, these results show that the human hepatocytes in LHM can impact the function of the hypothalamus, resulting in a substantial loss of rhythmicity or a change of rhythmicity of gene expression in the SCN and the ARC.

**Figure 5.**
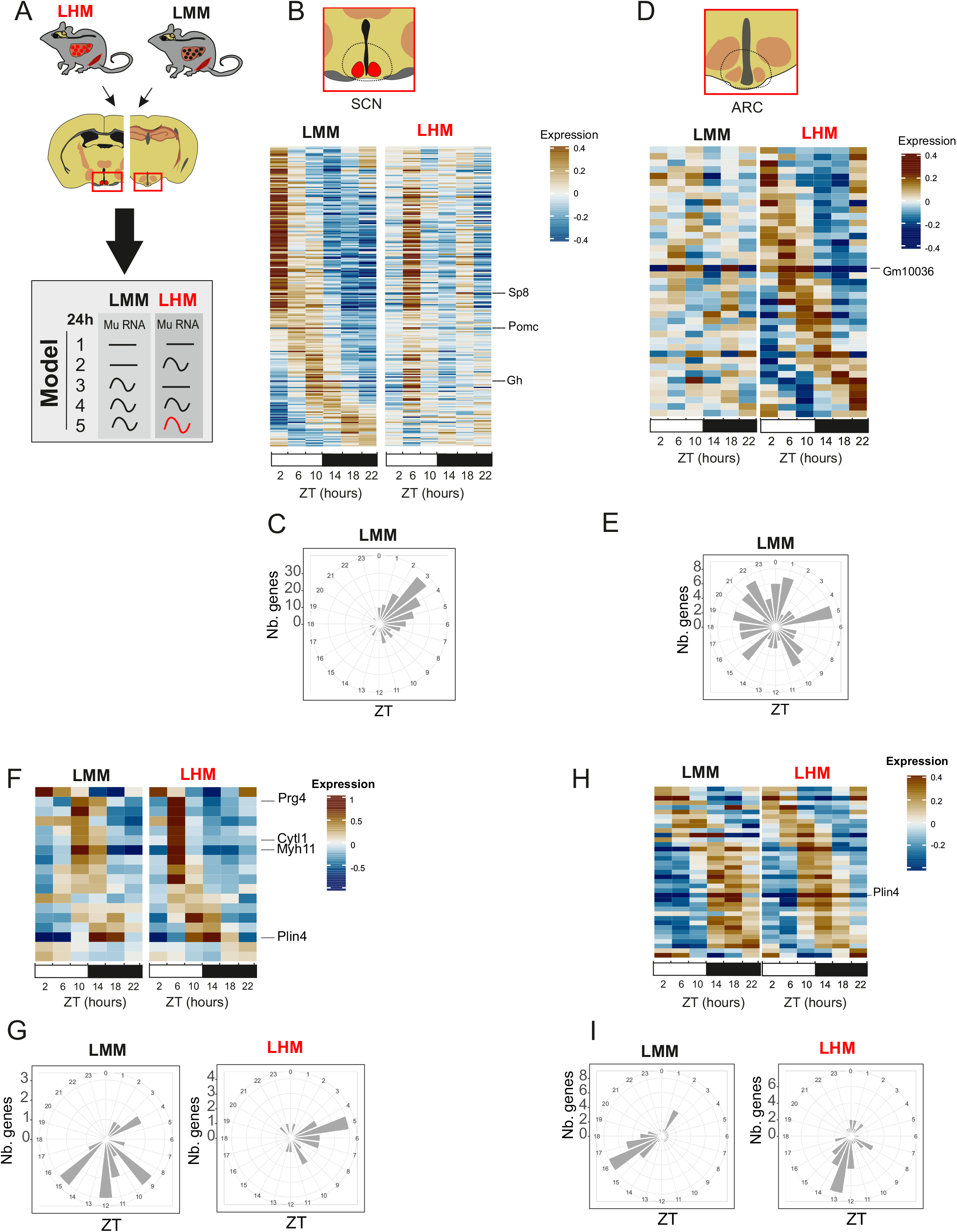
Engrafted human hepatocytes impact rhythmic gene expression in the hypothalamus. **A**) Experimental design for SCN and ARC samples collection prior to RNA extraction, sequencing and analyzing according to rhythmic properties, alteration of rhythmic gene expression of murine transcript in the liver of LMM (black) and LHM (red) assessed by model selection (model 1–5): black line, stable transcription; black wave, rhythmic transcription; red wave, rhythmic profiles with different rhythmic parameters (i.e., phase and/or amplitude). **B**-**E**) Heat maps of normalized rhythmic mRNA levels (BICW > 0.5, log2 amplitude > 0.5) and radial plot of the distribution of the peak phase of expression of the cycling genes and in the SCN (**B, C**) and ARC (**D, E**) in LMM (black) and LHM (red) from genes that lost rhythmicity in LHM (model 3). **F**-**I**) Heat maps of normalized rhythmic mRNA levels (BICW > 0.5, log2 amplitude > 0.5) and radial plot of the distribution of the peak phase of expression of the cycling genes and in the SCN (**F, G**) and ARC (**H, I**) in LMM (black) and LHM (red) from genes that show altered rhythmic parameters in LHM (model 5).

### Engraftment of human hepatocytes reveals the ability of hepatic signals to feedback on the central pacemaker

While the SCN is required for the coordination of feeding and drinking rhythms (Stephan and Zucker, 1972; Van den Pol and Powley, 1979), a functional SCN counteracts and delays the synchronization of mice to day feeding. Consequently, SCN-lesioned animals synchronized their circadian peripheral clocks more rapidly to day feeding (Saini et al., 2013; Sinturel et al., 2021). On the other hand, feeding rhythm is a very potent synchronizing signal for peripheral tissues and can uncouple peripheral oscillators from the central clock (Damiola et al., 2000; Stokkan et al., 2001). Because LHM show an advanced phase of physiology and a global loss of rhythmic gene expression in the SCN, we speculate that the engraftment of human hepatocyte can impact the capacity of the SCN to oppose the synchronization of animal physiology to feeding during the light phase.

To test this hypothesis, LMM and LHM were subjected to a 5-days measurement of baseline rhythmic physiology under *ad libitum* feeding, followed by a 7-days imposed feeding during the light phase before returning to *ad libitum* feeding (**Figure 6A**). As described above, we observed an advance in the phase of feeding and metabolic outputs at baseline conditions (**Figure 6B, 6E, and S6A**) which were abolished during an imposed feeding regimen (**Figure 6C, 6F, S6A**). Strikingly, upon day 5 of return to *ad libitum* feeding, the advanced phase in LHM reappeared (**Figure 6D, 6G, S6A**), showing that this phase advance is an intrinsic property of the SCN of LHM. While around 40% of the locomotor activity of LMM shift to the day, as expected as an adaptation to feeding during the light phase, they kept the circadian component of their locomotor activity at night, even after 6 days of this regimen (**Figure 6H, 6I, 6L and S6B**). However, LHM loose this circadian component and shift rapidly most of their daily locomotor activity to the light phase (**Figure 6H,6 I, 6L and S6B**). Of note, this lot of LMM and LHM did no significantly differ in body weight and body fat (**Figure S6C**), suggesting that the change in entrainment properties did not result from metabolic impairment. More strikingly, while the drinking activity of LMM, which is imposed by the central clock, is essentially restricted to the dark phase, it is shifted very rapidly to the light phase in LHM. Thus, approximately 90% of drinking activity during the light phase was observed already after 3 days (**Figure 6J, 6K, 6M and S6B**). This observation shows that the implantation of human hepatocyte modifies the entrainment properties of the SCN of LHM that becomes more sensitive to feeding cues and let them adapt more rapidly to daytime feeding paradigm, reminiscent of the adaptation observed in SCN lesioned animals (Saini et al., 2013).

**Figure 6.**
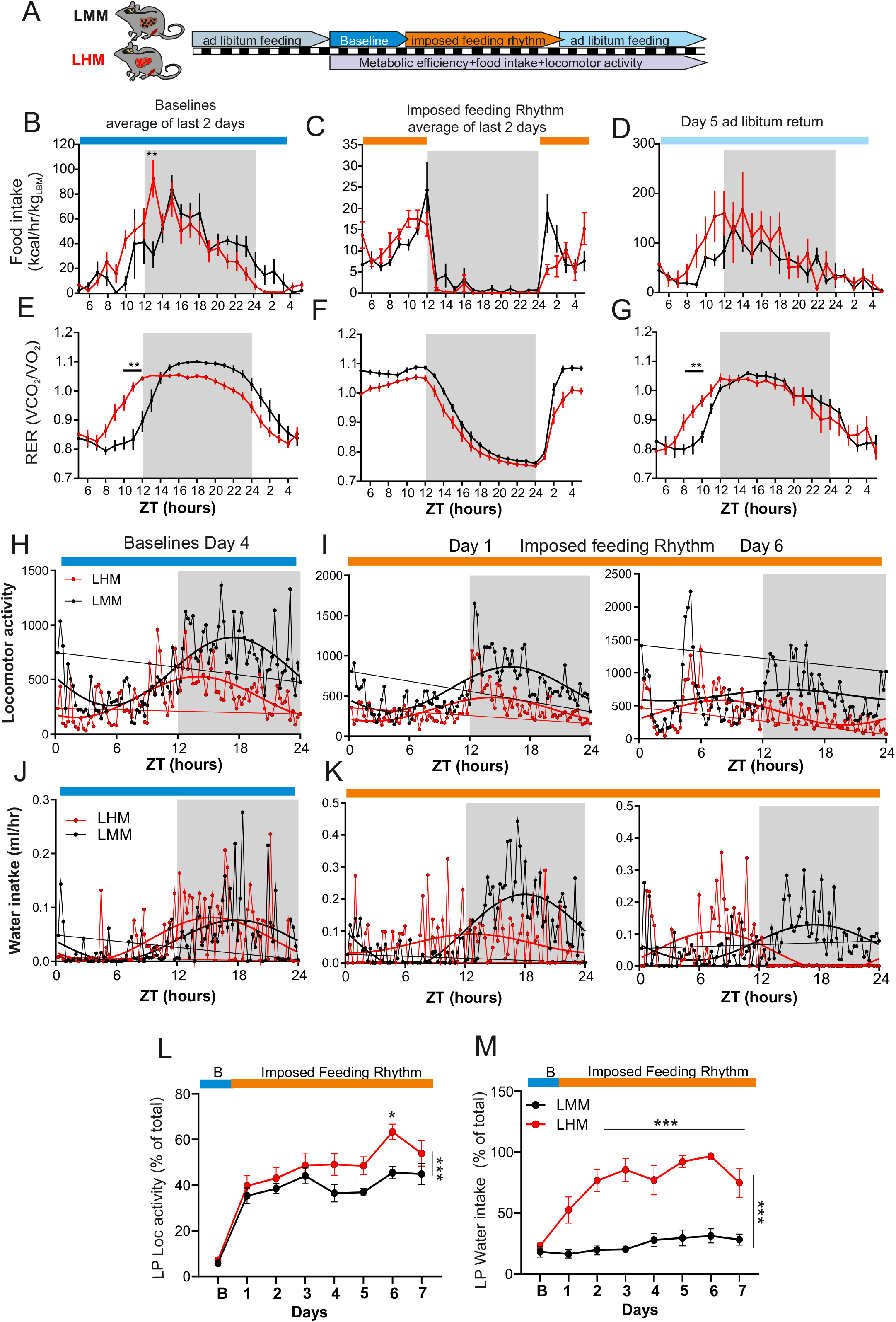
Engraftment of human hepatocytes reveals the ability of hepatic signals to feedback on the central pacemaker. **A**) Experimental design for the imposed feeding regimen during the light phase. **B**-**G**) 2-days average analysis of rhythmic food intake (**B-D**) and RER (**E**-**F**) during the baseline period (**B, E**), imposed daylight feeding (**C, F**), or day 5 after return to *ad libitum* feeding (**D, G**) in LMM (black) and LHM (red) (n=6 per groups). **H**-**K**) Representative of average distribution and non-linear cosinor fitting of baseline day 4 (**H, J**) and after 1 and 6 days of imposed light phase feeding rhythm (**I, K**) for locomotor activity (**H, I**) and water consumption (**J, K**). **L, M**) Evolution locomotor activity (**L**) and water intake (**M**) during the light-phase (LP) after transition from baseline (blue) to imposed feeding during the light phase (orange) as a percentage of baseline value. Data are expressed as mean +/-SEM (n=6 animal per condition). * P<0.05, ** P<0.01, *** P<0.0001.

## DISCUSSION

The data presented here shows that the engraftment of human hepatocytes can impact on the diurnal physiology and behavior of the grafted mice. Strikingly, the human hepatocytes not only advance the phase of surrounding mouse liver cells, but also the phase of distant muscle cells, likely through the change of global rhythmic physiology and behavior. Even more surprising is the discovery that the human hepatocytes also influence the rhythmic property of hypothalamic centers, probably by impacting the rhythmic gene expression in the SCN and the ARC. These effects resulted in a shorter circadian period, a phase advanced activity under a light-dark cycle, including the feeding rhythm, and a faster synchronization to feeding cues. This suggests that a circadian oscillator in peripheral organs with different circadian properties can feedback on brain centers to control circadian physiology and behavior. While keeping central and peripheral clock in phase is clearly of paramount importance, there are still some uncertainties as for the relative independence or total subordination of peripheral oscillators versus the central clock. Uncertainties also exist on how a peripheral organ rhythm can possibly modulate or even entrain the central clock. Although the SCN is required to maintain synchrony between the peripheral organs (Izumo et al., 2014; Sinturel et al., 2021; Yoo et al., 2004), the hepatocyte oscillator can work autonomously (Koronowski et al., 2019) and the synchrony between the hepatocytes persists in SCN-lesioned mice (Izumo et al., 2014; Sinturel et al., 2021). However, while the hepatocyte clock has the ability to modulate the circadian physiology of other liver cells or distant organs (Guan et al., 2020; Manella et al., 2021), no impact on the central circadian clock has been reported. Hence, our data show for the first time that the hierarchy between SCN and hepatocytes in the control of the circadian physiology can be challenged since peripheral oscillators with different circadian rhythms and responses to systemic cues impact the central oscillator.

Several mechanisms could be involved. The liver-brain axis could encompass rhythmic secretion of liver-borne molecules acting centrally and/or peripherally together with nervous pathway connecting these two organs. However, while several food-related hormones or metabolites modulate the synchronization of peripheral clocks or feeding behavior in response to nutrient signals, none of them have been shown to impact the central circadian clock and modulate the circadian period of the animals (Bookout et al., 2013; Chen et al., 2019; Landgraf et al., 2015). Of importance, the hepatocytes in LMM and LHM result from the migration of injected hepatocytes in the spleen and transfer to the liver, followed with positive selection of human hepatocytes and of healthy murine hepatocytes (Azuma et al., 2007; Ellis et al., 2013; Minniti et al., 2020). In this procedure LMM and LHM retain the architecture of liver-brain vagal connection. Hence, it is possible that the nervous connection between the liver and the brain may have routed hepatic information able to modify the central control of circadian physiology. In turn, modified central control will impede on peripheral organs function through the hypothalamic-pituitary-adrenal axis and/or the autonomic nervous system, resulting in peripheral tissue synchronization.

The misalignment of central and peripheral clocks due to shift work, jet lag, or desynchrony between food and light signals (abnormal feeding pattern) has been implicated in various diseases including sleep disorders, psychiatric disorders, and cardiometabolic disease, including atherosclerosis, obesity, diabetes, and non-alcoholic fatty liver disease (Bishehsari et al., 2016; Mokhlesi et al., 2019; Shan et al., 2018; Tan et al., 2018; Vetter et al., 2018). Conversely, the metabolic syndrome is associated with disruption of circadian rhythms and sleep (Lee et al., 2017; Zimmet et al., 2019). Mice exposed to a high fat diet exhibit an alteration of their circadian clocks and feeding behavior (Kohsaka et al., 2007) and circadian desynchrony impacting phase coherence between peripheral organs and communication between peripheral and central clocks (Dyar et al., 2018). Moreover, high fat diet induces an advance of phase of the liver circadian clock (Pendergast et al., 2013) and alters the response of the central clock to light signals (Mendoza et al., 2008). Likely these changes are rather a consequence of the metabolic syndrome caused by this regimen than the high fat diet *per se*. Indeed, female or male mice from inbred lines that did not develop metabolic syndrome did not display alteration of their circadian and feeding behavior (Buckley et al., 2021; Palmisano et al., 2017). Interestingly, liver diseases in human were shown to alter the function of the peripheral and central circadian clocks. Liver cirrhosis was associated with sleep disturbances, delayed phase of diurnal skin temperature cycle (Bruyneel and Serste, 2018; Cordoba et al., 1998; Garrido et al., 2017), and shifted plasma melatonin and cortisol profiles, two hormonal phase markers of the human SCN clock (Montagnese et al., 2010). In conclusion, our study provides the first evidence that the implantation of hepatocytes with independent self-sustained circadian clock with different responses to systemic cues can advance behavioral and metabolic rhythms through action on hypothalamic centers.

While many studies in rodent have underscored the implication of hepatocyte circadian clock on animal physiology, we describe here in mice with hepatocyte humanization the ability of peripheral cells to feedback to the central clock to the extent of redefining the free-running period and the phase of behavior and metabolism. Our study illustrates the benefit of chimeric animals to decrypt the impact of peripheral tissues in the control of circadian physiology. By taking advantages on the species-specific difference in cellular physiology, LHM allow to probe hierarchical organization of the circadian system which would otherwise be inaccessible in a homogenous rodent or human context. While our study does not provide a simple molecular mechanism by which human hepatocyte are capable of entraining tissues, our results force us to reconsider the importance of peripheral cues as Zeitgebers for behavior and physiology. The reciprocal relationship between central and peripheral clocks should be taken into account in future studies aimed at the dissection of circadian physiology and metabolism. We believe that our results beg to reconsider the importance of the hepatic signals as Zeitgebers. In that view, it is formally possible that the cardiometabolic diseases associated with disrupted circadian rhythms could primarily originate in peripheral tissues which in turn will promote desynchrony in the other tissues.

## EXPERIMENTAL PROCEDURES

### Animal ethics

All animal experiments were performed with approval of the Animal Care Committee of the Université Paris Cité (CEB-25-2016; number: 004667.02) and the veterinary office of the Canton of Vaud, Switzerland (authorization VD 3170). Chimeric animals were generated at Yecuris Corporation (Tualatin, OR, USA) or Karolinska Institutet (Stockholm, Sweden). This study was approved the animal care committee and complies with the Declaration of Helsinki, and ethical approvals (2010/678-31/3 and S82-13) were obtained from local authorities. All animal work was conducted according to approved Institutional Animal Care and Use Committee (IACUC, Yecuris Corporation) protocol DN000024 and NIH OLAW assurance #A4664-01. The protocols follow the NIH Guide for the Care and Use of Laboratory Animals.

### Animals

*Fah*^-/-^, *Rag2*^-/-^, *Il2rg*^-/-^ mice (FRG) were crossed with non-obese diabetic (NOD) mouse strain to create FRGN mice (Yecuris,Tualatin, OR, USA), whose livers can be fully repopulated with human or murine hepatocytes. Human liver tissue and hepatocytes were obtained through the Liver Tissue Cell Distribution System, and the studies were exempted by IRB 0411142 since no human subjects were involved (University of Pittsburgh). Human and murine hepatocytes were transplanted via splenic injections that allowed to repopulate the mouse liver by maintaining on 2-(2-nitro-4-fluoromethylbenzoyl) cyclohexane -1,3-dione (CuRx™ Nitisinone (NTBC) in the drinking water, a medication that prevents hepatocyte death when FAH deficiency is present. Engraftment is sustained over the life of the animal with an appropriate regimen of Nitisinone (20-0026) and prophylactic treatment of Sulfamethoxazole (SMX) / Trimethoprin (TMP) antibiotics (20-0037). Due to the impaired immune system of these stains, they are susceptible to respiratory infections. Mice received prophylactic antibiotic treatment cycles in order to minimize respiratory infections. Treatments with a combination of SMX/TMP for 5 days twice a month greatly reduces the occurrence of common respiratory infections as well as subsequent secondary infections. The final concentration of SMX/TMP drinking water is 640µg/ml, 128 µg/ml (TMP) (CuRx™ SMX/TMP Antibiotic Cat# 20-0037, Yecuris corporation) and 3% Dextrose (Sigma-Aldrich ref D9434). The homozygous deletion of fah in FRGN mice which produces a fumaryloacetoacetate dehydrogenase deficiency that disrupts tyrosine metabolism, leading to buildup of hepatotoxic metabolite, fumarylacetoacetate. With the administration of 2-(2-nitro-4-fluoromethylbenzoyl) cyclohexane -1,3-dione (CuRx™ Nitisinone (NTBC) Cat# 20-0026), the intracellular accumulation of furmarylacetoacetate is blocked and the mice live normal immune deficient mouse lifespans. Mice are cycling on 8mg/l NTBC treatment with SMX/TMP antibiotic and 3% of Dextrose for 3 days every 5 weeks. To maintain hydration of the mice during the extended period off Nitisinone, it is best to use sterile drinking water that contains 3% Dextrose. Because the dextrose provides an opportunistic environment for bacterial growth, new containers of sterile dextrose water need to be prepared every week as described previously (Azuma et al., 2007). Chimeric animals were generated at Yecuris Corporation (Tualatin, OR, USA). Ten weeks old male mice FRGN® repopulated with human hepatocytes (LHH, Hu-FRGN®) and FRGN® repopulated with murine hepatocytes (LMM, Mu-FRGN®) (Cat#10-0013 and Cat#10-0009, YECURIS corporation, Oregon, USA). Mice were maintained on on Pico Lab High Energy Mouse Diet 5LJ5 (LabDiet). Only mice with human hepatocyte repopulation >70% (corresponding to circulating levels of human albumin >3.5 mg/mL) were used in this study.

Detailed engraftment for experimental group is as follow: experimental group presented in **Figure 1, 2, Figure S1A, C, D, E, H, Figure S2** is composed of 12 mice transplanted with human female donor F1 HHF13022 (n=12, LHM-F1) and 12 mice transplanted with NOD murine donor (n=12, LMM-F1). Experimental group presented **Figure 3, S1F, S3** is composed of 5 mice transplanted with human male donor F1 HHM30017 and 10 mice transplanted with human female donor HHF13022 (n=15, LHM-F1) and 12 mice transplanted with NOD murine donor (n=12, LMM-F1). Experimental group presented **Figure 4 A-H, S1B, S4** is composed of 6 mice transplanted with human female donor F1 HHF17006 plus 2 mice transplanted with human male donor F1 (n=12, LHM-F1) and 12 mice transplanted with NOD murine donor (n=12, LMM-F1), experimental group in Figure 4 I-K is composed of 5 mice transplanted with human male donor F1 HHM30017 and 10 mice transplanted with human female donor HHF13022 (n=15, LHM-F1) and 12 mice transplanted with NOD murine donor (n=12, LMM-F1). Experimental group presented **Figure 5, 6, S1G, S5, S6** is composed of 12 mice transplanted with human female donor HHF13022 (n=12, LHM-F1) and 12 mice transplanted with NOD murine donor (n=12, LMM-F1).

### Time-resolved tissue collection

Except otherwise noticed, mice were kept under diurnal lighting conditions (12-hr light 12-hr dark) with unrestricted access to food. Mice were sacrificed by at 4-hour intervals over 24 hours corresponding to ZT2, ZT6, ZT10, ZT14, ZT18 and ZT22 by decapitation and tissues were dissected, snap-frozen in liquid nitrogen, and stored at -80°C until further processing.

### Serum samples

Blood samples were collected in serum collection tubes and allowed to clot for 1 h at 4 °C on ice. Tubes were centrifuged at 10,000 g for 5 min at 4 °C and serum was stored immediately at -80 °C. During the data collection, serum samples were aliquoted and thawed no more than twice.

### Human albumin measurement

After 1,000 or 10,000x dilution with Tris-buffered saline, human albumin concentration was measured with the Human Albumin ELISA Quantitation Kit (Bethyl E80-129) according to the manufacturer’s protocol, to monitor the progress of humanization of transplanted mice. 1000 μg/mL of circulating human albumin correlates with ∼20% engraftment of human cells, 2000 μg/mL with ∼40% and mice with 4000 μg/mL of human albumin showed approximately at ∼80% of human cells in the repopulated liver (Ellis et al., 2013).

### Plasma rodent growth hormone and IGF-1 measure

Plasma rodent growth hormone (Ref.22-GHOMS-E01) and rodent IGF-1 (Ref. 22-IG1MS-E01) content of each sample was measured in duplicate with a colorimetric assay provided by ALPCO corporation. The absorbance from each sample was measured in duplicate using a spectrophotometric microplate reader. The intra- and inter-assay coefficients of variation for these kits were around 0.3-8%. Plasma samples were tested in duplicate within one assay, and the results were expressed in terms of the standards supplied (ng/mL).

### Quantification of plasma lipoprotein lipids

Lipoproteins were separated from 2.5 μL of individual plasma samples by size exclusion chromatography (SEC), using a Superose 6 PC 3.2/300 column (GE Healthcare Bio-Sciences AB, Uppsala, Sweden). Lipoproteins were eluted as a fraction appearing in the exclusion volume of the sepharose column that contained VLDL, then LDL and last HDL. TG, concentrations were calculated after integration of the individual chromatograms (Parini et al., 2006; Pedrelli et al., 2014), generated by the enzymatic-colorimetric reaction with the respective following kits, TG GPO-PAP (Roche Diagnostics, Mannheim, Germany).

### Indirect calorimetry metabolic analysis

The indirect calorimetry system carries out preclinical non-invasive and fully automated measurement of food and water intake, O_2_ consumption, CO_2_ production, respiratory quotient (as an indicator of glycolic or oxidative metabolic status), whole energy expenditure together with tridimensional (X, Y, Z) spontaneous activity and fine movement (Phenomaster, TSE Systems GmbH) as described (Joly-Amado et al., 2012). Mice were monitored for whole energy expenditure (EE) or Heat (H), oxygen consumption and carbon dioxide production, respiratory exchange rate (RER = VCO2/VO2, where V is a volume), and locomotor activity using calorimetric cages with bedding, food, and water (Labmaster, TSE Systems GmbH,

Bad Homburg, Germany). Ratio of gases was determined through an indirect open circuit calorimeter. This system monitors O2 and CO2 concentration by volume at the inlet ports of a tide cage through which a known flow of air is being ventilated (0.4 L/min) and compared regularly to a reference empty cage. For optimum analysis, the flow rate was adjusted according to the animal body weights to set the differential in the composition of the expired gases between 0.4 and 0.9% (Labmaster, TSE Systems GmbH, Bad Homburg, Germany). The flow was previously calibrated with O2 and CO2 mixture of known concentrations (Air Liquide, S.A. France). Oxygen consumption, carbon dioxide production and energy expenditure were recorded every 15 min for each animal during the entire experiment. Whole energy expenditure was calculated using the Weir equation respiratory gas exchange measurements. Food consumption was measured as the instrument combines a set of highly sensitive feeding and drinking sensors for automated online measurements. Unless noted otherwise mice had free access to food and water ad libitum. To allow measurement of every ambulatory movement, each cage was embedded in a frame with an infrared light beam-based activity monitoring system with online measurement at 100 Hz. The sensors for gases and detection of movement operated efficiently in both light and dark phases, allowing continuous recording. Mice were monitored for body weight and composition at the entry and the exit of the experiment. Body mass composition (lean tissue mass, fat mass, free water and total water content) was analyzed using an Echo Medical systems’ EchoMRI (Whole Body Composition Analyzers, EchoMRI, Houston, USA), according to manufacturer’s instructions. Briefly, mice were weighed before they were put in a mouse holder and inserted in MRI analyser. Readings of body composition were given within 1 min. Data analysis was performed on Excel XP using extracted raw value of VO2 consumed, VCO2 production (express in mL/h), and energy expenditure (kcal/h). Subsequently, each value was expressed by total body weight extracted from the EchoMRI analysis

### Imposed day feeding regimen

Mice were individually housed for at least 2 weeks and body weight monitored daily. Mice were then housed in cages allowing for measure of metabolic efficiency, food intake, water intake, and locomotor activity (Phenomaster system described previously). After a 10 days *ad libitum* feeding and baseline recording, mice were imposed a strict daylight feeding for 8 days (food available from ZT0 to ZT12) and return to normal regimen for 8 days.

### RNA extraction and sequencing

Liver RNA were isolated using the RNeasy kit from QIAGEN according to the manufacturer’s recommendations. The integrity and the concentration of each RNA sample was assessed using Agilent 2100 bioanalyzer. cDNA was prepared following the NEBNext® Ultra™ II Directional RNA Library Prep Kit protocol (New England Biolabs). The libraries were sequenced using paired-end 150 bp sequencing on a HiSeq 4000 System (Illumina). Muscle, ARC, and SCN RNA were isolated using the RNAdvance Tissue kit from Beckman Coulter according to the manufacturer’s recommendations. The integrity and the concentration of each RNA sample was assessed using Agilent Fragment Analyzer-96 and with Quant-It Ribogreen from Life Technologies. Libraries were prepared following the TruSeq Stranded mRNA Kit protocol from Illumina and sequenced using paired-end 126 bp sequencing on a HiSeq 2500 System (Illumina).

### Analyse of rhythmicity

STAR 2.4.0i (Dobin et al., 2013) was used to map RNA-seq reads onto a chimeric genome consisting of the Mus musculus (GRCm38/mm10) and Homo sapiens (GRCh38/hg19) reference sequences and to quantify the number of uniquely mapped reads per gene and organism. For the liver data, human and mouse genes with on average more than 10 read counts in LMM or LHM were retained. Human and mouse genes were matched using mouse-human orthologs database from Ensembl. To assess differential rhythmicity and mean differences of gene expression in RNA-Seq raw count data, we used the *dryR* method based on a model selection framework with generalized linear models (Weger et al., 2021). Genes with a BICW (likelihood to belong to the chosen rhythmic model) larger than 0.3 (liver) or 0.5 (muscle, SCN, ARC) and a log2 amplitude > 0.5 in at least one condition were used for the functional and gene set enrichment analysis

### Functional and gene set enrichment analysis

#### GO analysis

Gene set enrichment analysis was performed using enrichR (Kuleshov et al., 2016) with GO Biological Process (2021) or GO Molecular Function (2021) databases. Terms with adjusted p-value < 0.1, combined score > 50 and odds ratio > 6 were selected.

#### mTOR targets

Supplementary table with differential expression analysis was downloaded from (Bai et al., 2021). Genes were considered as differentially expressed between wildtype and respectively *Raptor* KO and *Tsc1* KO mice at ZT12, if absolute log2 fold change was larger than 1 and p-value smaller than 0.05. Statistical overrepresentation of those genes sets in genes upregulated (log2FC > 1, *dryR* chosen model mean 2) or downregulated (log2FC < 1 and *dryR* chosen model mean 2) in LHM v LMM liver was computed using a hypergeometric test.

#### System-driven genes

Supplementary table for rhythmicity analysis was download from (Koronowski et al., 2019). For each category (WT, *Bmal1* KO, *Bmal1* Liver-RE), genes were defined as rhythmic if the p-value was smaller than 0.05. Statistical overrepresentation of those gene sets in the different rhythmic models of LHM and LMM (BICW > 0.5 and log2 amplitude > 0.5) was assessed using a hypergeometric test.

#### CLOCK-target genes

The CLOCK-target gene set was retrieved from Alhopuro *et al*. (Alhopuro et al., 2010). We used hypergeometric testing to calculate overrepresentation of the geneset within the statistical models identified by *dryR*. A threshold of BICW > 0.5 was applied to the data and for the liver dataset an additional threshold on amplitude log2FC > 0.5.

### Sex-biased gene expression and activation of the STAT5 pathway

To predict activities of transcription factors and sex biased genes, we employed ISMARA (Balwierz et al., 2014) on the count data after variance stabilizing transformation (vst) (Love et al., 2014). Target genes for sex biased STAT5 targets and sex biased genes in mouse liver were taken from Zhang *et al*. ((Zhang et al., 2012)) and Weger *et al*. (Weger et al., 2019), respectively.

### Detection of phospho-S6 and S6 ribosomal protein in mouse extracts

Liver from LMM and LHM were resuspended into lysis buffer (PBS 150 mM NaCl, pH 7,5, 1 % Triton X-100, 2 mM NaF, 2 mM Na_3_VO_4_, protease inhibitors), sonicated (10 sec, 10 % power) and lysed for 30 min at 4°C under agitation. The lysate was centrifuged for 20 min at 15,000 g (4°C) and the sample protein concentration was determined using Bradford’s method. 60 µg of each sample were separated by SDS-PAGE (4-12 %, Invitrogen) followed by a transfer onto a nitrocellulose membrane (0.22 µm, GE Healthcare) for 1 h at 4°C. A ponceau staining of the membrane was done to ensure equal protein loading. Membranes were then blocked in 5 % bovine serum albumin (BSA) in TBST (Tris 20 mM, 150 mM NaCl, pH 7.6, 0.1 % Tween-20) for 1 h and incubated overnight with phospho-S6 ribosomal protein antibody (#2211, Cell Signaling) in 1 % BSA TBST at 4 °C. The next day, membranes were washed 3 times with PBST prior to incubation with secondary antibody for 1 h at room temperature. Membranes were then washed 3 more times and the signal was detected by chemiluminescence using ECL Prime reagent (GE Healthcare) on an Amersham Imager 600 detection system (GE Healthcare, France). Membranes were later stripped using stripping buffer (Tris 60 mM, pH 6,8, 2 % SDS, 100 mM β-mercaptoethanol) for 30 min at room temperature, blocked in 5 % BSA TBST for 1 h and reprobed with S6 ribosomal protein antibody (#2317, Cell Signaling) using the same protocol as above. Signal analysis were performed using ImageJ software taking as standard reference a fixed threshold of fluorescence.

### Statistical analysis

All statistical comparisons were performed with Prism 9 (GraphPad Software, La Jolla, CA, USA). All the data were analyzed using either Student t-test (paired or unpaired) with equal variances or One-way ANOVA or Two-way ANOVA. Cosinor parameters (acrophase, mean, period, amplitude) results were analyzed by simple comparisons (unpaired t-test). In all cases, significance threshold was automatically set at p < 0.05. ANOVA analyses were followed by Bonferroni *post hoc* test for specific comparisons only when overall ANOVA revealed a significant difference (at least p < 0.05).

### Data availability

RNA-Seq raw files have been deposited in NCBI’s Gene Expression Omnibus (GSE212079) https://www.ncbi.nlm.nih.gov/geo/query/acc.cgi?acc=GSE212079

## ACKNOWLEDGMENTS

We would also like to thank all of the consortium members of the EU-funded research project HUMAN (Health and the Understanding of Metabolism, Aging and Nutrition, http://www.fp7human.eu/) who have directly and indirectly contributed to the discussion of the results. We thank Olja Kacanski for administrative support, Isabelle Le Parco, Ludovic Maingault, Angélique Dauvin, Aurélie Djemat, Florianne Michel, Magguy Boa, and Daniel Quintas for animals’ care and Sabria Allithi for genotyping. We are grateful to Professor Ueli Schibler for critical input in the manuscript. We acknowledge the technical platform Functional and Physiological Exploration platform (FPE) of the Université de Paris (BFA - UMR 8251), the animal core facility Buffon of the Université de Paris/Institut Jacques Monod, and Aymen Kaabi (Nestlé Research) for animal experiments.

## AUTHOR CONTRIBUTIONS

**Conceptualization**: A.S.D., M.Q., P.P., F.G., and S.L.; **Investigation**: A.S.D., M.Q., C.G., J.C., R.G.P.D., J.B., B.D.W., A.C., S.M., J.P., M.K., L.L.V., M.E.M., M.P., F.G.; **Formal Data Analysis**: C.G., B.D.W., E.C.; **Resources:** J.B., E.M.W.; **Funding acquisition**: P.P., F.G., S.L.; **Supervision**: A.S.D., P.P., F.G., S.L.; **Writing – original draft**: F.G., S.L.; **Writing – review & editing**: all authors.

## FUNDING

This work has been financed by the HUMAN project (grant agreement no. 602757) within the EU’s Seventh Framework Program (FP7) for research, technological development, and demonstration. F.G. receives support from the Institute for Molecular Bioscience, The University of Queensland. M.Q was supported by a postdoctoral contract from the Galician Government type A (Xunta de Galicia ED481B2014/039-0) and type B (Xunta de Galicia ED481B2018/004). M.Q, is currently funded by a research contract “Miguel Servet” (CP21/00108) from the ISCIII and co-funded by the European Union. We acknowledge funding supports from the Centre National la Recherche Scientifique (CNRS) and the Université Paris Cité.

## CONFLICT OF INTEREST

J.B. was formerly President and Chief Executive Officer of Yecuris Corporation. E.M.W. was formerly Senior Scientist at Yecuris Corporation. C.G., B.D.W., A.P., S.M., J.P., M.K., and F.G. are or were employees of Société des Produits Nestlé SA.

## SUPPLEMENTARY FIGURE LEGENDS

**Figure S1. Characterization of the humanized mouse model and its impact on liver gene expression**

**A**) Plasma level for human albumin in LHM mice 2 months after primary hepatocyte engraftment and prior to sacrifice at different ZT time. Human albumin is non-detectable in LMM.

**B**) Plasma triglyceride (TG) Lipoprotein profile and in LMM and LHM.

**C**) Proportion of human (red) and murine (black) reads counts in the liver of humanized (LHM) or “murinized” (LMM) model.

D) Number of genes in all models of rhythmic gene expression (Figure 1B) in the liver of LMM and LHM (BICW > 0.3, log2 amplitude > 0.5).

**E**) Enrichment of up- and down-regulated genes in the liver of LMM and LHM for genes up- and down-regulated in the liver of mice KO for the negative (*Tsc1* KO) or positive (*Raptor* KO) regulators of mTOR (see methods).

**F**) Western blot analysis of Phospho-S6 Ribosomal Protein (Ser240/244) and total S6 Ribosomal Protein and non-linear cosinor fitting of P-RPS6 signal normalized by ponceau signal. Analysis of rhythmicity done with *dryR* (BICW = 0.572 for rhythmicity only in LMM).

**G**) Circulating murine growth hormone (GH) and Insulin-like growth factor-1 (IGF-1) in LMM and LHM. Data are expressed as mean +/-SEM (n=9-11 animals for each condition). *** P<0.0001.

**H**) Top. Female-(left) and male(right)-biased gene expression from control and humanized mouse livers at indicated Zeitgeber Time (color). Bottom. The predicted activity of sex dependent (female (♀, left) and male (♂, right) target genes of STAT5 in the mouse liver. P values were determined by a two-way ANOVA (time, condition), P value for condition is reported (n = 12 mice per condition).

**Figure S2. Impact of the engrafted human hepatocytes on liver rhythmic gene expression**

**A, B**) Heat maps (**A**) and radial plot of the distribution of the peak phase (**B**) of normalized rhythmic mRNA levels (BICW > 0.5, log2 amplitude > 0.5) in the liver of LMM (black) and LHM (red). Genes are assigned to model 2 in which only human genes are rhythmic in LHM animals while non-rhythmic in LMM.

**C**) Enrichment of the rhythmic genes (-log10 p-value, hypergeometric test, see method) in the liver of LMM and LHM ranked in the 15 models of rhythmicity for genes rhythmic in all combinations of WT, *Bmal1* KO (KO), and hepatocyte specific *Bmal1* rescue in *Bmal1* KO animals (liver-RE). While genes directly regulated by BMAL1 (rhythmic in WT and liver-KO) are enriched in models 4 (rhythmic only in LMM), 11 (same rhythm in LMM and mouse and human transcripts in LMM), and 14 (rhythmic in all conditions but different phase in LMM), rhythmic genes dependent on systemic cues regulated by BMAL1 (rhythmic only in WT) are enriched only in model 4, showing that the rhythm of many of these genes depends on systemic signals regulated by the circadian clock in non-liver tissues.

**Figure S3. The muscle circadian clock of liver humanized mice is phase advanced**

**A**) Number of genes in all models of rhythmic gene expression (Figure 3A) in the muscle of LMM and LHM (BICW > 0.5, log2 amplitude > 0.5).

**B, C**) Heat maps (**B**) and radial plot of the distribution of the peak phase (**C**) of normalized rhythmic mRNA levels (BICW > 0.5, log2 amplitude > 0.5) in the muscle of LMM (black) and LHM (red). Genes are assigned to model 3 in which mRNA are only rhythmic in LMM.

**Figure S4. Human hepatocytes advance the phase of circadian metabolism and behavior of liver humanized mice**

**A**-**D**) Cosinor analysis of the rhythmic parameters mean level (a) and amplitude (b) for locomotor activity (**A**), food intake (**B**), RER (**C**), and fat oxidation (**D**). Data are expressed as mean +/-SEM (n=6 animals per condition). * P<0.05, ** P<0.01

**E**) Averaged values for body weight, lean body mass, and fat mass in LMM (black) and LHM (red) at the beginning of the experiment. Data are expressed as mean +/-SEM (n=6 per condition). P values were determined by a one-way ANOVA. * P<0.05, ** P<0.01.

**Figure S5. Engrafted human hepatocytes impact rhythmic gene expression in the hypothalamus A**) Expression level of the ARC-enriched *Agouti related transcript* (*Agrp*) and the SCN-enriched *Aralkylamine N-Acetyltransferase* (*Aanat*) in the dissected ARC and SCN of LMM (black) and LHM (red). **

**B, C**) Number of genes in all models of rhythmic gene expression (Figure 5A) in the SCN (**B**) and ARC (**C**) of LMM and LHM.

**D, E**) Rhythmic expression of the circadian clock regulated genes *Nr1d1* (D) and *Dbp* (E) in the SCN and ARC, respectively, of LMM and LHM.

**F**) Enrichment of the rhythmic genes in the SCN and ARC of LMM and LHM ranked in the 5 models of rhythmicity for genes bound by CLOCK in ChIP-Seq experiment (see Method).

**Figure S6. Engraftment of human hepatocytes reveals the ability of hepatic signals to feedback on the central pacemaker**

**A**) Cosinor analysis of the acrophase (c) for RER during baseline (left), imposed feeding during the light phase (middle) and after return to *ad libitum* feeding (right). Data are expressed as mean +/-SEM (n=6 per condition). ** P<0.01, *** P<0.0001.

**B**) Averaged values and kinetics of water intake in LMM mice (black upper panel) and LHM mice (red, lower panel) during baselines (blue, left Y axis) and imposed feeding rhythm (orange, right Y axis).

**C**) Averaged values for body weight, lean body mass and fat mass in LMM (black) and LHM (red) at the beginning of the experiment. Data are expressed as mean +/-SEM (n=6 per condition). P values were determined by a one-way ANOVA.

## Notes

https://www.ncbi.nlm.nih.gov/geo/query/acc.cgi?acc=GSE212079

