## Supplementary figures and images for "Mice with humanized livers reveal the involvement of hepatocyte circadian clocks in rhythmic behavior and physiology"

### Figure S1

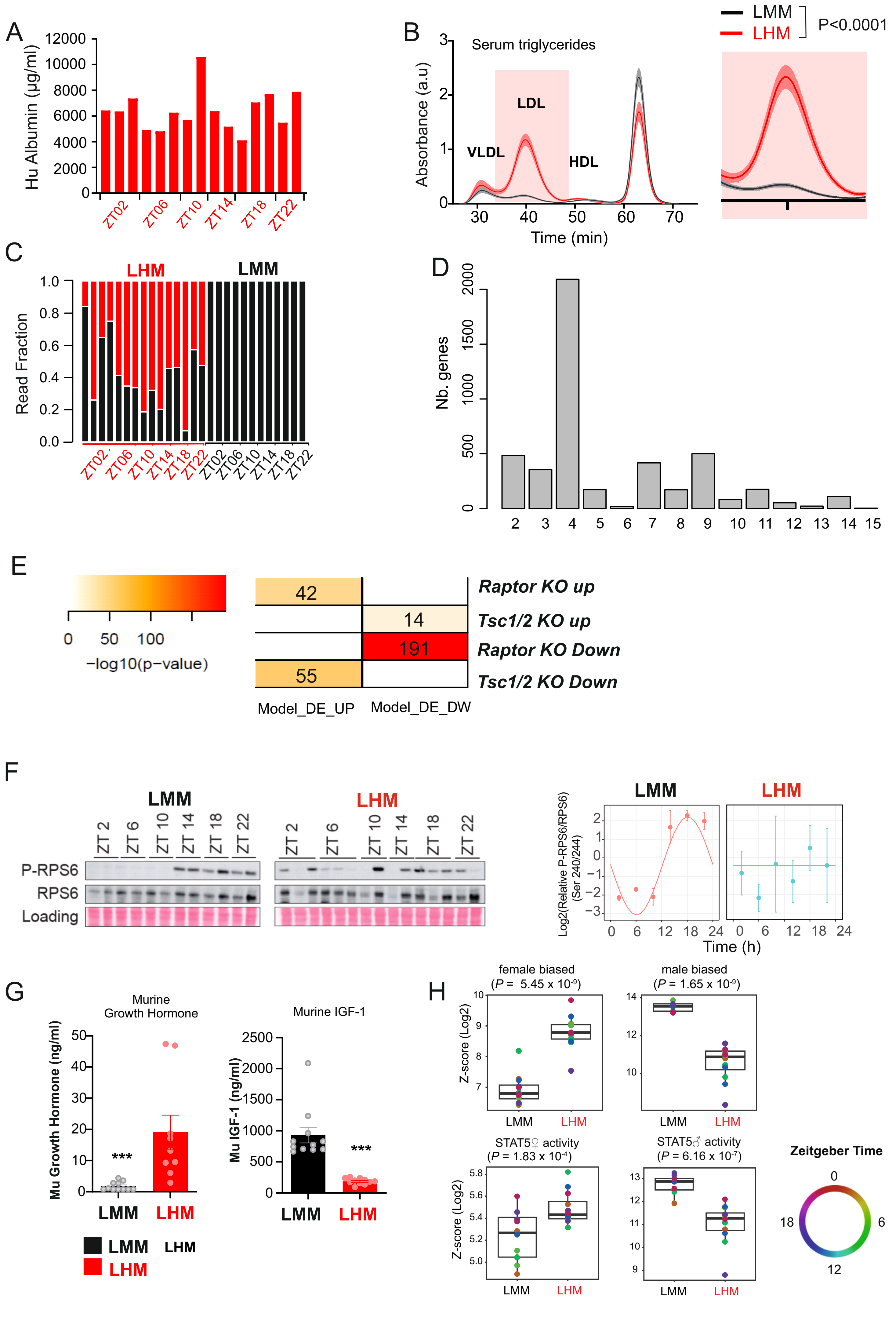

### Figure S2

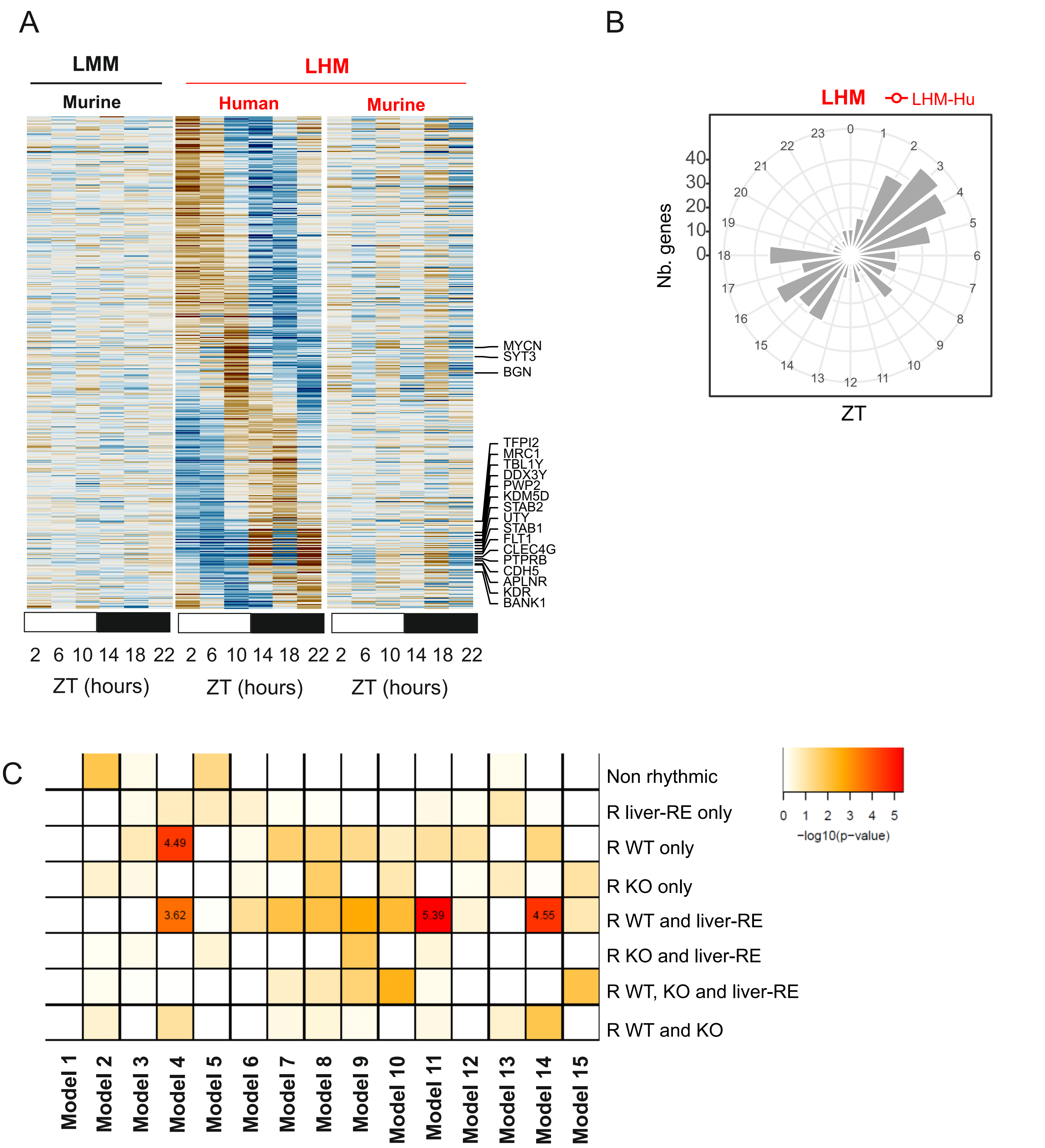

### Figure S3

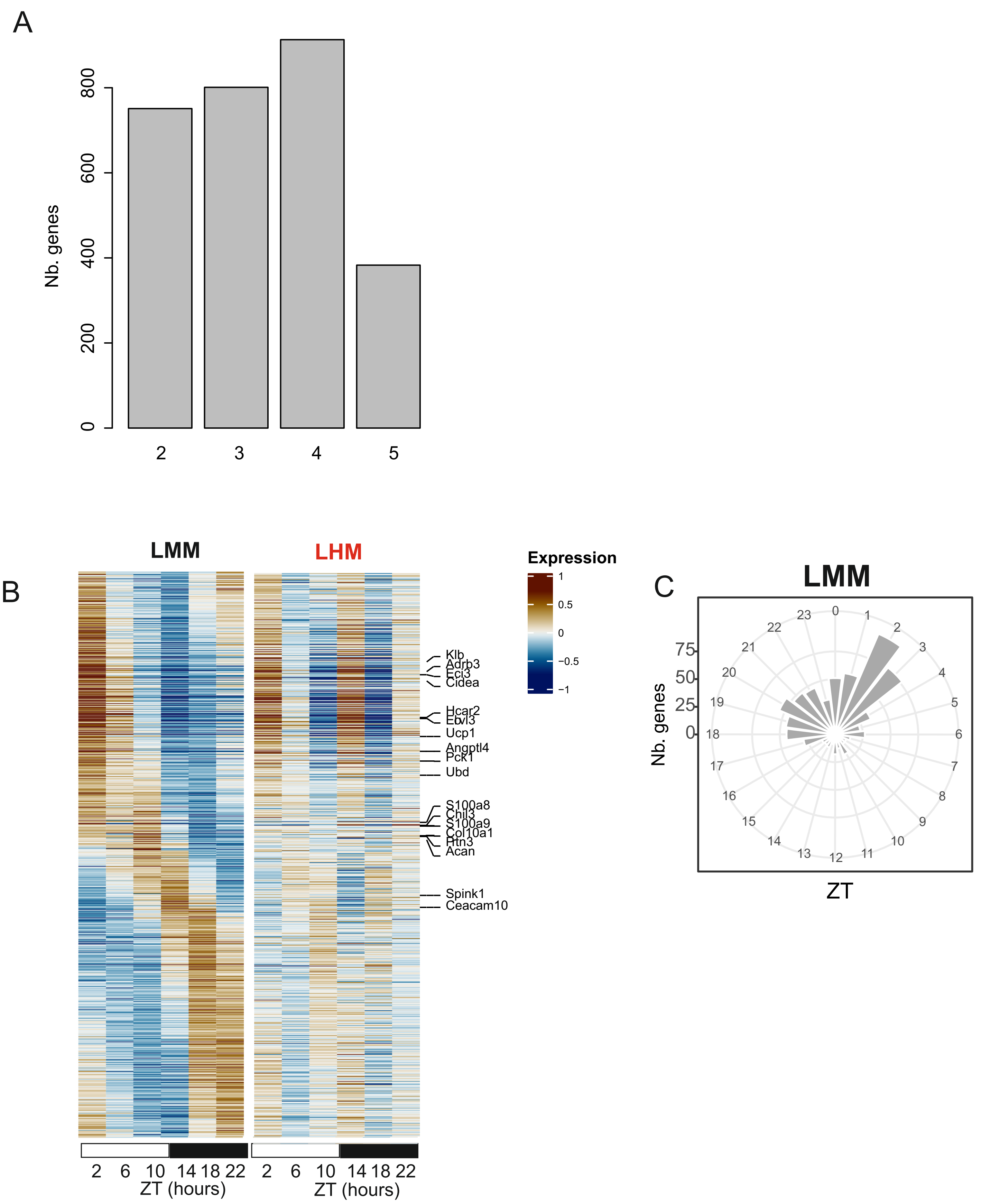

### Figure S4

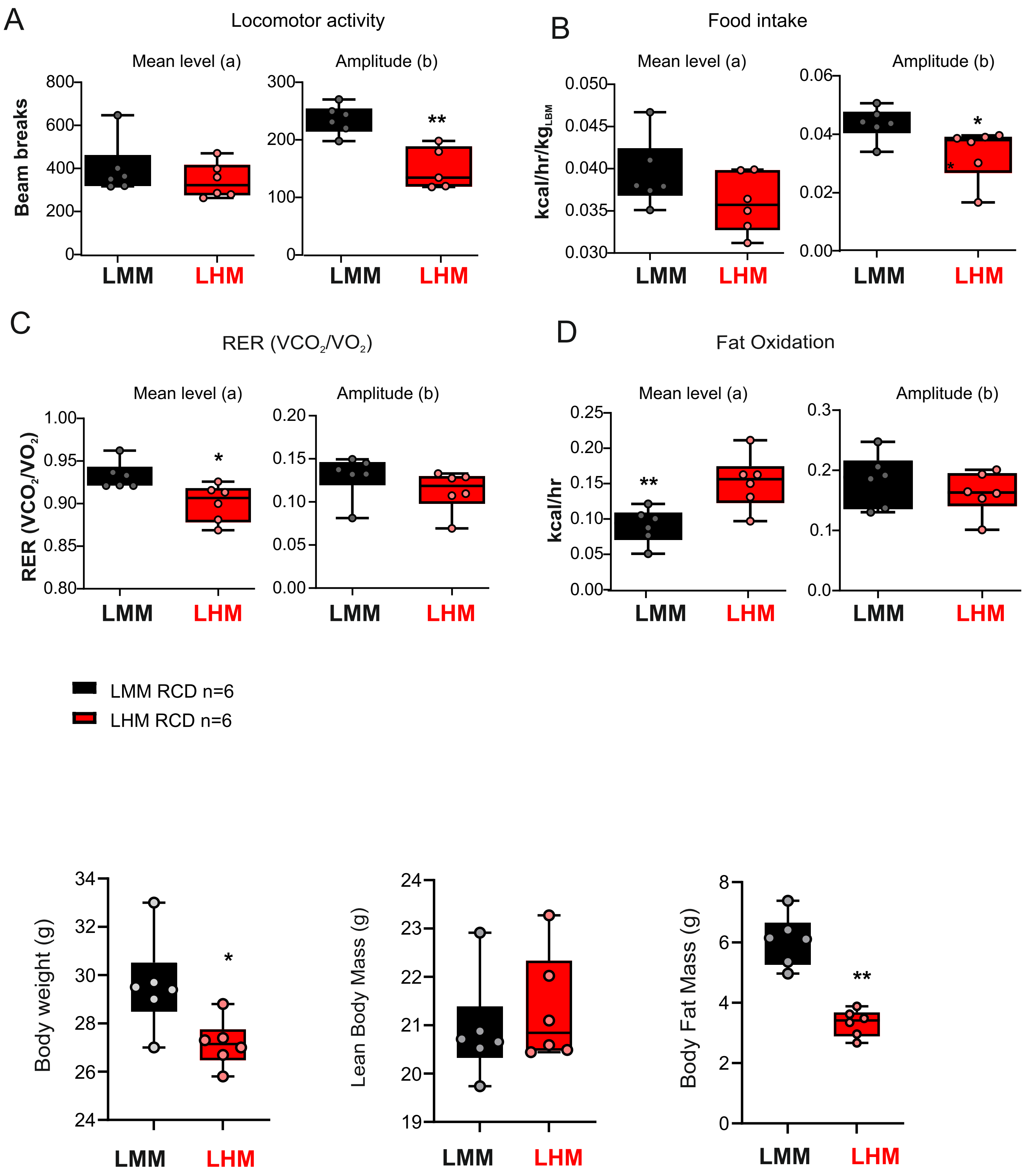

### Figure S5

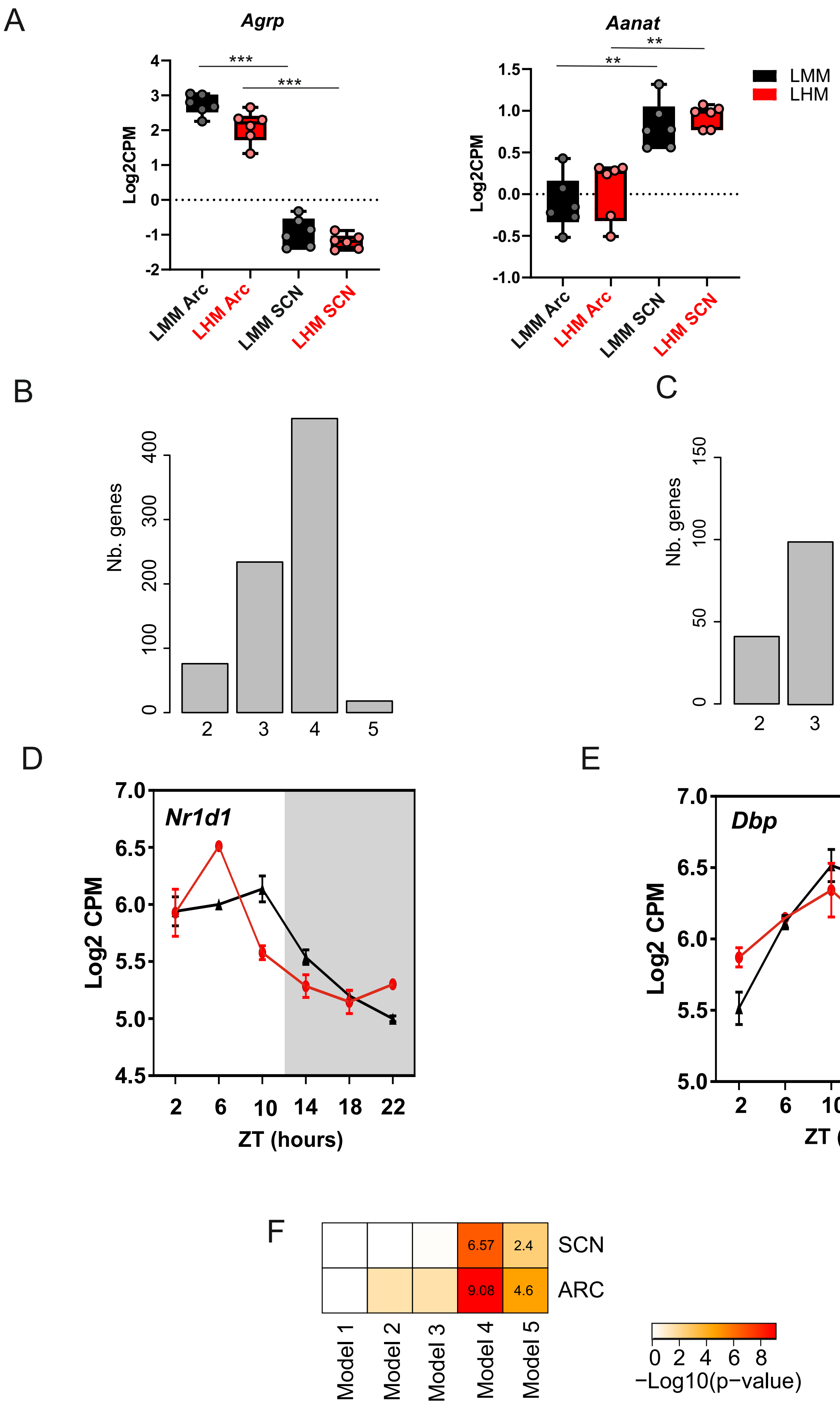

### Figure S6

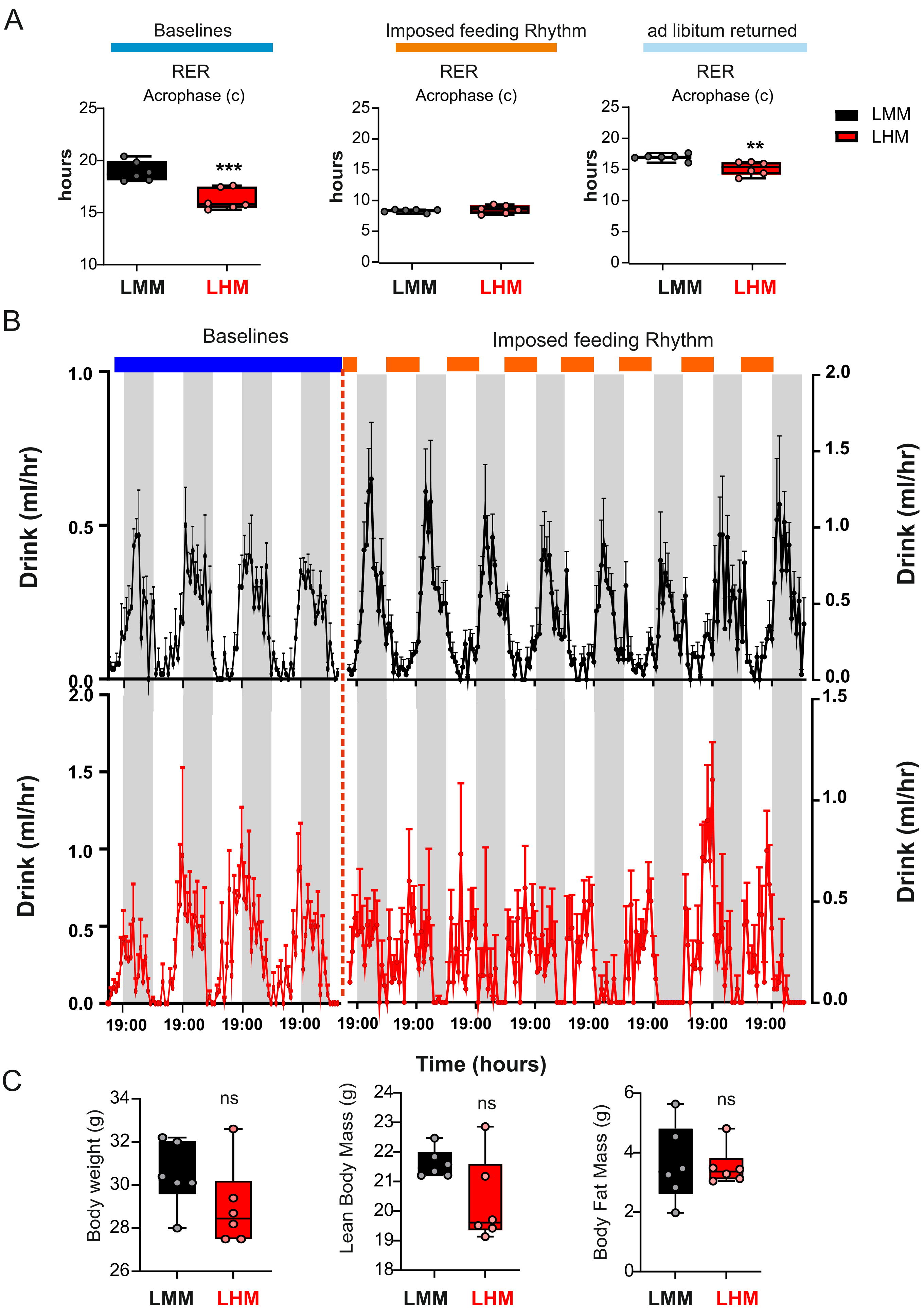
